# Comparing self- and hetero-metacognition in the absence of verbal communication

**DOI:** 10.1101/585240

**Authors:** Laurène Vuillaume, Jean-Rémy Martin, Jérôme Sackur, Axel Cleeremans

## Abstract

The ability to infer how confident other people are in their decisions is crucial for regulating social interactions. It is unclear whether one can read others’ confidence in absence of verbal communication and whether one can infer it as accurately as for one’s own confidence. To address these questions, we used an auditory task in which participants had to guess the confidence of someone else performing the task or to judge their own confidence in different conditions (i.e., while performing the task themselves or while watching themselves performing the task on a pre-recorded video). Results show that participants are able to guess the confidence of other people as accurately as when judging their own. Crucially, we show that hetero-metacognition is a flexible mechanism relying on different cues according to the context. Our results support the idea that metacognition leverages the same inference mechanisms as involved in theory of mind.

## INTRODUCTION

Metacognition — ‘cognition about cognition’ — is typically characterized as involving two distinct but interconnected processes: evaluation and control. Metacognitive evaluation involves monitoring the quality of first-order processing, such as memory, perception, language, reasoning and so on (e.g. Beran et al., 2012; Dienes & Perner, 2002; Hampton, 2001; Koriat, 2000; 2007; 2012; Nelson & Naren, 1990; Proust, 2007; Schwartz, 1994; Smith et al., 2003). Then, based on such metacognitive evaluation, people can deploy different control strategies in order to regulate their cognitive activities. As an illustration, having spent some time studying her lesson, a student may judge that she is still in trouble when rehearsing it and may decide to continue studying, as a result.

Research has shown that a large part of metacognitive evaluation or monitoring can take place outside conscious awareness, calling for a further distinction between implicit versus explicit metacognition (Shea et al., 2014; for similar distinctions see e.g., Koriat, 2000, 2007; Dokic & Martin, 2012; 2015; 2017; Martin & Dokic, 2013). Shea et al., (2014) conceptualise this distinction within a ‘dual systems’ view of metacognition: System 1 metacognition — implicit metacognition — is shared with other species (e.g., Summerfield & Yeung, 2013) and refers to processes that use metacognitive representations (i.e., representations about object-level representations) to control behaviour in an automatic way, without the subject being conscious of such representations. System 2 metacognition — explicit metacognition — on the other hand, might be uniquely human (Shea et al. 2014) and refers to the conscious use by the subject of metacognitive representations, leading to non-automatic voluntary monitoring and regulating control strategies.

Recent theorizing often assumes that much of cognitive control takes place outside System 2 (see, for instance, Shea et al., 2014). Thus, cognitive control mechanisms can be triggered by visually masked, unconscious stimuli (van Gaal et al., 2008, 2009, Sumner et al., 2007). But, if implicit metacognition is doing much of the work, what is the function of explicit, or System 2, metacognition? It has been proposed (Shea et al., 2014; Frith, 2012) that System 1 metacognition is for the control of intra-personal cognitive activities while System 2 metacognition, is for supra-personal cognitive control. In other words, System 2 metacognition has the computational resources of broadcasting intra-personal metacognitive information to others and, therefore, allows for the regulation of group behaviour (e.g., to coordinate sensorimotor activities between members of a group executing a common task). In the context of joint perceptual decision making, evidence shows that people indeed communicate their metacognitive representations, namely their confidence in their perceptual decisions, and that, under certain conditions, the communication of such metacognitive information leads to improved joint perceptual decisions (Bahrami, Olsen, Latham et al., 2010) — this is the “two-heads-better-than-one” effect (Bahrami et al., 2010; Koriat, 2012). Importantly, communication or sharing of confidence is necessary for such joint perceptual decision benefits to occur even in the presence of external feedback about the accuracy of the perceptual decision of both subjects of the dyad (conversely, the presence of external feedback is not necessary when confidence is shared). The improving effect of informational exchange between members of a team is not limited to perceptual discrimination and has been shown to improve problem solving (Cooper & Kagel, 2005) or reasoning (Maciejovsky et al., 2013), for instance.

Other theoretical proposals (Baumeister & Masicampo, 2010; Masicampo & Baumeister, 2013) share with the above the hypothesis that explicit metacognition, and conscious thoughts more generally, aims at regulating interpersonal cognitive control. In addition, in a recent computational account of confidence judgements in one’s first-order performances, Fleming and Daw (2017) proposed that intra-personal confidence judgements are generated in a way similar to evaluating others’ confidence in their own performance. Thus, many approaches converge on the notion that voluntary, explicit metacognition evolved as a means to improve social functioning.

This perspective presupposes that one has the cognitive resources to read others’ confidence. However, people’s ability to read others people’s confidence has so far received little attention. This stands in contrast with substantial research dedicated to the theory of mind ability to read others’ emotions (e.g., Zhou, Majka & Epley, 2017). Of course, in many situations it is just a matter of verbal communication: subject A says to subject B how uncertain she is about such or such event. In many other situations, verbal communication cannot be carried out as easily or even trusted. Imagine for instance that you are competing with someone or playing poker. While neither of you wants to share information, it can nevertheless be crucial to read the other’s confidence. In other daily life settings such as a romantic date or a job interview, one may not be able to rely as much on verbal communication as on other cues. Likewise, teachers need be able to carry out online evaluations about whether their students are keeping up with the pace.

Here, we investigated three main questions: First, are people able to read others’ confidence in the absence of verbal communication and, if it is indeed the case, what are the cues through which this is accomplished? In a significant paper, Patel et al. (2012) have shown that people are indeed able to read other people’s confidence in a visual discrimination task through the simple observation of the kinematics of others’ actions. Participants were shown two intervals that contained six Gabors arranged in a circular fashion around a fixation point. All the Gabors but one had the same contrast and participants had to decide which interval comprised the “oddball” stimulus. Participants took their decisions in displacing a marble into one of two holes corresponding to the first and the second interval. By means of different sensors, kinematics of decision-related actions were recorded. The observation task consisted in observing the video-recorded hands of anonymous participants performing the task. In addition, Patel et al., (2012) have demonstrated that the ability to read others’ confidence from the kinematics of their actions is based on one’s own movement kinematics properties when executing the task oneself. Patel et al. (2012) thus suggest that reading others confidence rests upon motor simulation mechanisms. Here, we surmised that movement cues are not the only ones people may use when assessing someone else’s confidence, especially when the participant observing the other has also access to the stimuli. Hence, we made the paradigm more ecologically valid by having people observe actual confederates executing the task.

The second question we investigated is to what extent performing the task oneself gives one privileged access in assessing confidence when compared to observing someone else performing the task. In other words, is there a first-person perspective benefit when assessing confidence, or does assessing one’s own confidence leverage exactly the same machinery as that involved when assessing someone else’s confidence (Carruthers, 2009; Fleming & Daw, 2017)?

The third question we explored is whether one has a privilege at inferring the confidence of an observed participant when the participant is oneself (by means of a video recording) *versus* someone else, when stimuli are not available. One could indeed conjecture that the link between task difficulty and confidence is so strong that potential first-person perspective cues are overridden by this link.

To explore these questions we designed an auditory pitch discrimination task in which participants had to decide which of two pure tones presented successively had the higher pitch — a first-order decision — and to rate their confidence in their response — a second-order decision (Figure 1). Pairs of participants were tested together. In one condition, participants performed the task separately (Baseline condition, Figure 1A); in another condition, while one participant was performing the task, the other was observing her doing it and had access to the stimuli (Full-Observation condition, Figure 1B). In what follows the term ‘observer’ denotes the participant observing the other participant performing the task, whom we will call the ‘agent. In the Full-Observation condition, the observer was to judge the confidence of the agent on each trial (of course, the confidence ratings of the agent were hidden from the observer). In the Partial-Observation condition (Figure 1C), the observer was also to judge the confidence of the agent but he did not have access to the stimuli. Finally, in the Self-Observation condition (Figure 1D), each participant observed herself doing the task from a video recording of their Baseline condition but did not have access to the stimuli themselves.

**Figure 1.**
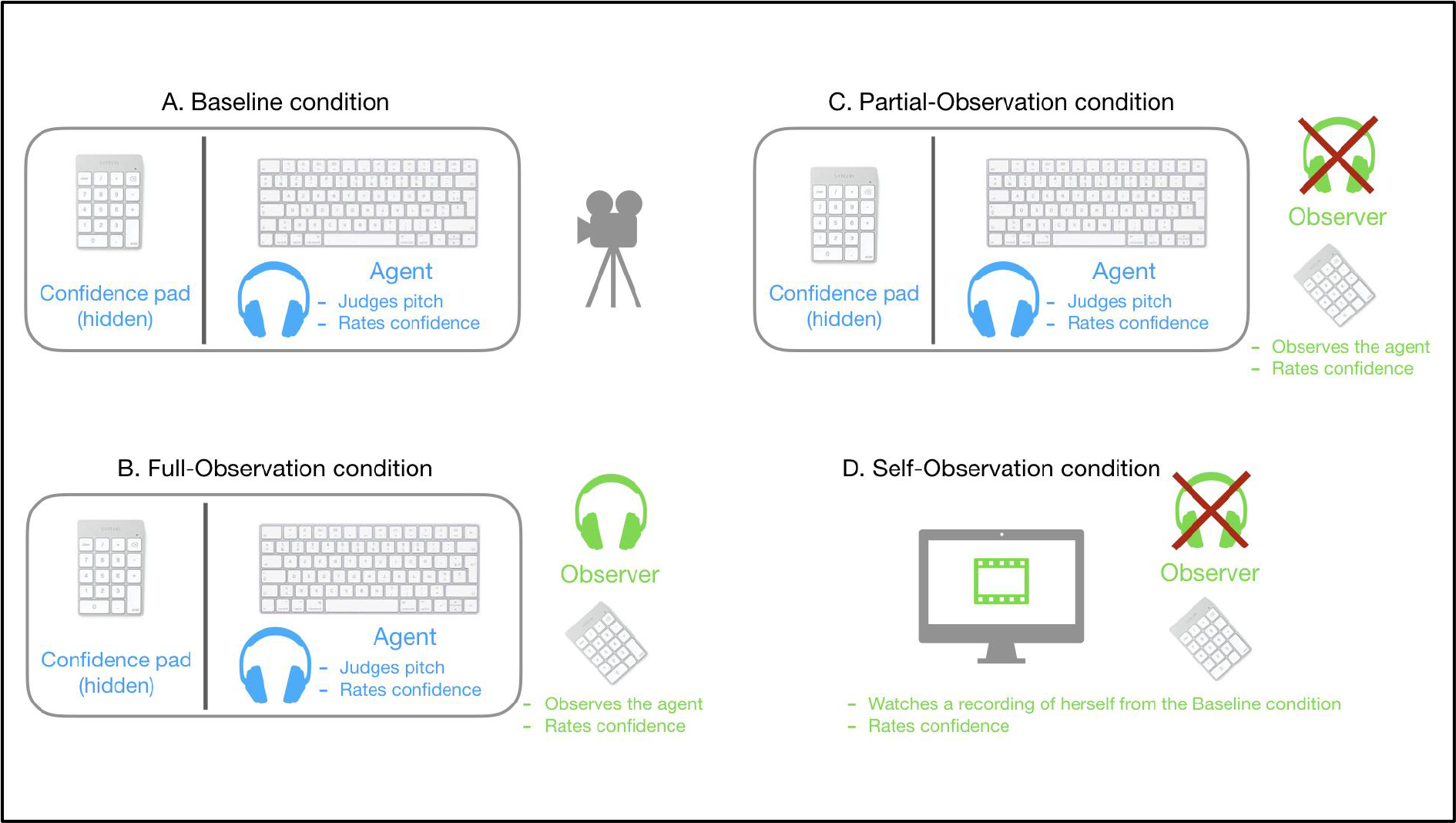
Experimental design. A. Baseline condition, B. Full-Observation condition, C. Partial-Observation condition and D. Self-Observation condition.

We hypothesised that participants would be able to judge the confidence of the agent and that it would be easier for observers to judge agents’ confidence in the Full-Observation condition than in the Partial-Observation condition because of the strong cue that task difficulty constitutes in confidence rating. However, if reading others’ confidence in absence of verbal communication is indeed possible, the performance of observers should also be positive in the Partial-Observation condition. We additionally hypothesised that, in this condition, agents’ response times might be a strong cue for observers. Indeed, it has been shown that response times are an important cue used to infer one’s own confidence (Kiani, Corthell & Shadlen, 2014; Desender, Van Opstal & Van den Bussche, 2017). Furthermore, if there is a kind of first-person perspective benefit in assessing confidence, people should be better in evaluating their own confidence in the Baseline condition than in evaluating the confidence of someone else in the Full-Observation condition. Finally, with the same reasoning, people should be better in inferring the confidence of an observed participant when the participant is herself (Self-Observation condition) *versus* someone else (Partial-Observation condition), in absence of the stimuli.

## RESULTS

### Agent performance at the first- and second-order level

First, regarding the first-order task (i.e., pitch discrimination task), a repeated-measure ANOVA showed no effect of condition on type 1 sensitivity (*d’*) (F(1.10, 18.64) = 1.01, p > 0.3, η_p_^2^ = 0.06) or criterion (F(1.22, 20.79) = 0.35, p > 0.4, η_p_^2^ = 0.02). However, a repeated-measures ANOVA showed differences in mean response times across the different conditions (F(1.39, 23.71) = 14.34, p < 10^−3^, η_p_^2^ = 0.60). Specifically, paired *t*-tests indicated that response times were shorter in the Full-Observation (Mean = 1.41 s, SD = 0.26) and Partial-Observation (Mean = 1.35 s, SD = 0.28) conditions than in Baseline condition (Mean = 1.97 s, SD = 0.53) (Full-Observation vs. Baseline: p < 10^−2^, Partial-Observation vs. Baseline: p < 10^−3^), but there was no difference between the Partial- and Full-Observation condition (p = 0.10, BF = 0.32).

Second, with regards to the second-order task, we found no effect of condition on confidence ratings (F(1.06, 18.05) = 0.65, p > 0.4, η_p_^2^ = 0.04) or confidence ratings variability using standard deviation as a measure of variance (F(1.24, 21.15) = 0.26, p > 0.4, η_p_^2^ = 0.02). This indicates that the first-order performance and confidence estimates of the agent were not impacted by the different conditions.

Third, we compared the metacognitive sensitivity of the agent across the different conditions using A_ROC_ as a measure of this type 2 sensitivity. A repeated-measure ANOVA showed no difference between conditions (F(1.51, 25.66) = 1.24, p = 0.30, η_p_^2^ = 0.07).

### Observer mean confidence level across conditions

The mean confidence level of the observer did not differ between conditions (F(1.74, 29.64) = 2.80, p = 0.08, η_p_^2^ = 0.14) nor did the confidence variability (F(1.70, 28.85) = 2.70, p = 0.09, η_p_^2^ = 0.14).

### Observer ability to read agent confidence

Before comparing the relation between the observer and the agent confidence between the different conditions, we checked whether this relation was significant within each condition. For each condition, we fitted a linear mixed-effects model of the confidence of the observer, with confidence of the agent as fixed effect and intercept for participants as random effect. Results confirm that the observer was indeed capable of judging the confidence of the agent in each condition (Full-Observation: estimate = 0.26, t = 18.63, p < 10^−3^; Partial-Observation: estimate = 0.16, t = 11.94, p < 10^−3^; self-observation: estimate = 0.18, t = 13.97, p < 10^−3^).

Then, we compared the relations between the agent’s and observer’s confidences between these different conditions. We fitted a linear mixed-effects model of the confidence of the observer, with confidence of the agent and condition (Full-Observation, Partial-Observation and Self-Observation) as fixed effects and intercept for participants as random effect.

The first row in **Table 1** (intercept) estimates the average confidence of the observer in the Full-Observation condition for the lowest scale rating of the confidence of the agent. The observer had a significantly higher confidence than the agent when the latter reported guessing (estimate = 2.15, t = 21.50, p < 10^−3^).

**Table 1.** Regression coefficients for the linear mixed-effects model of the confidence of the observer in the three conditions.

|  | Estimate | SE | T | p |
| --- | --- | --- | --- | --- |
| Intercept | 2.15 | 0.10 | 21.50 | <0.001 |
| Confidence of the doer | 0.26 | 0.01 | 20.79 | <0.001 |
| Partial-Observation condition | 0.27 | 0.05 | 5.27 | <0.001 |
| Self-Observation condition | 0.01 | 0.05 | 0.23 | 0.82 |
| Confidence of the doer: Partial-Observation condition | -0.12 | 0.02 | -6.95 | <0.001 |
| Confidence of the doer: Self-Observation condition | -0.05 | 0.02 | -2.76 | <0.01 |

The second row estimates the regression slope between the confidence of the observer and the confidence of the agent in the Full-Observation condition, and shows that this relation is statistically significant (estimate = 0.26, t = 20.79, p < 10^−3^), indicating that the observer can track the confidence of the agent.

The third and fourth row of the model show that the confidence of the observer for the lowest scale rating of the agent was not significantly different between the Self-Observation condition and Full-Observation condition (estimate = 0.01, t = 0.23, p > 0.4), but was significantly higher in the Partial-Observation compared to the Full-Observation condition (estimate = 0.27, t = 5.27, p < 10^−3^).

Crucially, the fifth and the sixth row indicate that the relation between the confidence of the observer and the confidence of the agent was smaller in the Partial-Observation compared to the Full-Observation condition (estimate = − 0.12, t = − 6.95, p < 10^−3^) and in the Self-Observation compared to the Full-Observation condition (estimate = − 0.05, t = − 2.76, p < 10^−2^). This indicates a decrease in the capacity of the observer to adapt her confidence to the confidence of the agent both in the Partial-Observation and the Self-Observation condition compared to the Full-Observation condition (Figure 2).

**Figure 2.**
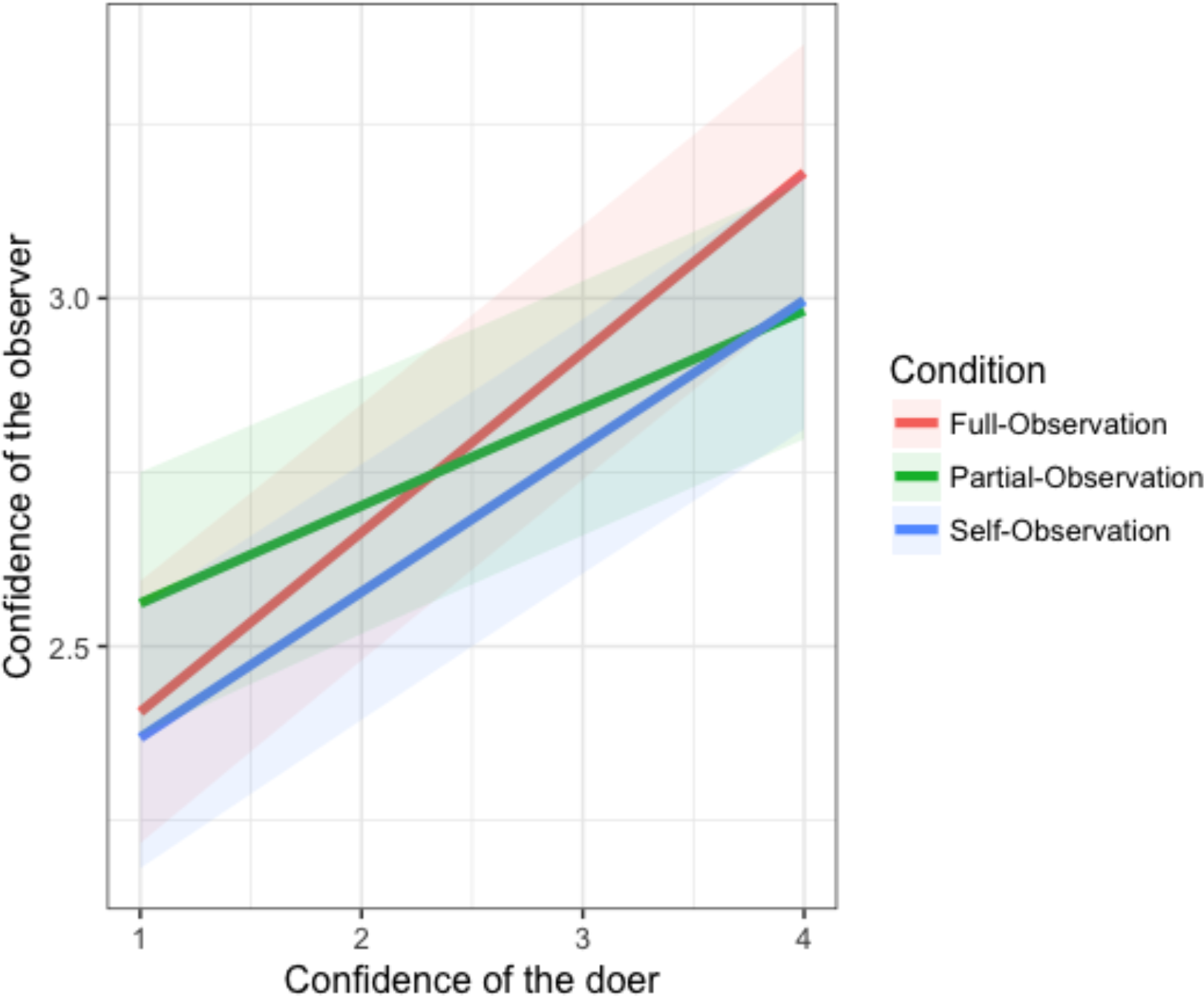
Regression slopes from the linear mixed-effects model between the confidence of the observer and the confidence of the agent in the Full-Observation (red), Partial-Observation (green) and Self-Observation condition (blue). The shaded area around each fit represents the 95% confidence interval.

Importantly, the difference between the Full- and Partial-Observation condition cannot be explained by differences stemming from the agent, as there was no difference in sensitivity, criterion, confidence or response times for the agent between these two conditions. In addition, it cannot be explained by differences in the confidence of the observer as it did not differ significantly between these two conditions.

Another linear mixed-effects model comparing only the Self-Observation condition (in which participants were judging their own performances in the baseline condition by means of video recording) to the Partial-Observation condition revealed significantly different intercepts (estimate = − 0.24, t = − 4.60, p < 10^−3^) and regression slopes (estimate = 0.06, t = 3.53, p < 10^−3^), indicating that the relation between the confidence of the observer and the confidence of the agent was stronger in the Self-Observation condition than in the Partial-Observation condition (note that in the Self-Observation condition the agent and the observer are actually the same subject).

### Do observers read agents’ confidence from their response times

We then thought to investigate which cues the observer relied on to judge the confidence of the agent. To do so, we explored whether and to which extent the confidence of the observer tracked the response times of the agent in the first-order task. The observer could indeed watch the speed with which the agent replied to the first-order task and use this information to answer on the confidence scale.

We used causal mediation analyses to test whether the effect of the confidence of the agent on the confidence of the observer was mediated by the response times of the agent (Figure 3). In each condition, a mediator mixed model was first fitted to predict the response times of the agent by the agent’s confidence. Then, an outcome mixed model was fitted to predict the confidence of the observer by the response times and the confidence of the agent. The mediation analysis was performed with these two models (using the mediation package; Tingley, Yamamoto, Hirose, Keele, & Imai, 2014) in order to test whether the influence of the confidence of the agent on the confidence of the observer was mediated by the response times of the agent.

**Figure 3.**
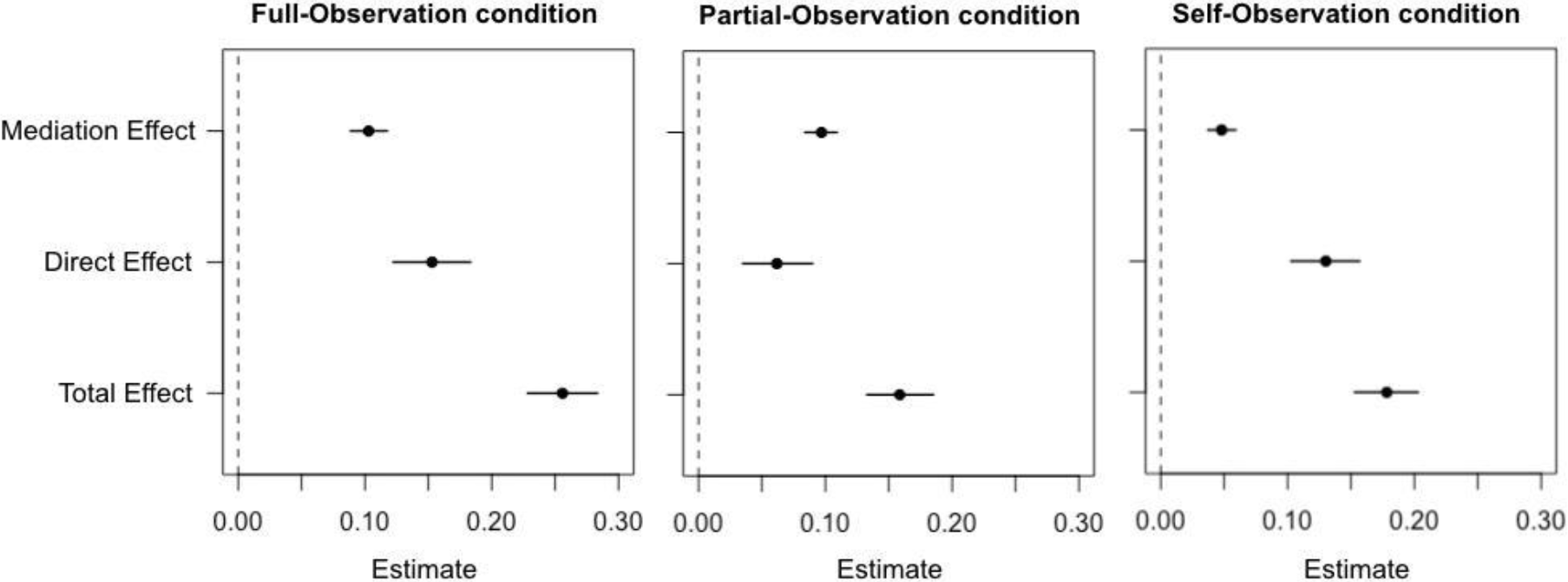
Results of the causal mediation analyses between the mediation and the outcome mixed regression models predicting the influence of the confidence of the agent on the confidence of the observer through response times. Error bars reflect quasi-Bayesian 95% confidence intervals.

In the Full-Observation condition the mediation analysis showed that from the total effect of the confidence of the agent on the confidence of the observer (β = 0.256, 95% CI = [0.228, 0.283], p < .001), there was 40.1% that was mediated by the response times of the agent (β = 0.103, 95% CI = [0.089, 0.118], p < .001). In the Partial-Observation condition, from the total effect of the confidence of the agent on the confidence of the observer (β = 0.159, 95% CI = [0.133, 0.185], p < .001), there was 61.1% that was mediated by the response times of the agent (β = 0.097, 95% CI = [0.084, 0.109], p < .001). In the Self-Observation condition, from the total effect of the confidence of the agent on the confidence of the observer (β = 0.178, 95% CI = [0.153, 0.203], p < .001), there was 26.9% that was mediated by the response times of the agent (β = 0.048, 95% CI = [0.037, 0.059], p < .001).

These findings suggest that response times of the agent were a crucial mediator between confidence of the agent and confidence of the observer in the Partial-Observation condition in which participants did not have access to the stimuli. However, this mediation was partly reduced in the Full-Observation condition and considerably reduced in the Self-Observation condition, suggesting that the observer relied less on the response times of the agent to estimate their confidence in this condition.

### Type-II signal detection theory

So far we have shown that participants are able to evaluate the confidence in others, even when they do not have access to the stimuli the agent is judging. In addition, mediation analyses indicate that to judge confidence in others, participants may use the agent response times, especially when they do not have access to the stimuli. Finally, regression analyses show a difference between the Partial- and Self-Observation conditions, with a stronger relation between the confidence of the agent and the confidence of the observer in the latter than in the former. This suggests that we do have some kind of privileged access to our own confidence. However, the cues the cognitive system is using are different between conditions as mediation analyses show that in the Self-Observation condition response times mediate to a lesser extent the relation between the confidence of the agent and the confidence of the observer. To further corroborate the results of regression analyses between the confidence of the agent and the confidence of the observer, we performed type-II signal detection theory analyses (SDT) (Maniscalco & Lau, 2014). Type-II SDT allows to compute the metacognitive sensitivity of individuals, that is their ability to discriminate between their correct and incorrect first-order responses. Here, we reasoned that if participants are able to read others’ confidence they should be able to discriminate between the correct and incorrect first-order responses of the agent, at least to some extent. So we computed A_ROC_ based on the confidence responses given by the observer and the first-order responses of the agent (which corresponds to the same subject in the Baseline condition).

As expected, a one-way one sample t-test showed that the A_ROC_ of participants judging themselves in the Baseline condition were significantly higher than 0.5 (p < 10^−6^). In the Full-, Partial- and Self-Observation conditions the A_ROC_ were also significantly higher than 0.5 (Full-Observation condition: p < 10^−4^; Partial-Observation condition: p < 0.01, Self-Observation condition: p = 0.03), suggesting that the metacognitive ability of the observer regarding the agent (or herself through video recording in the Self-Observation condition) was also higher than chance.

Analysis of variance revealed a significant difference between conditions (F(2.35, 39.99) = 8.08, p < 10^−3^, η_p_^2^ = 0.32), with higher A_ROC_ in the Baseline condition compared to the Partial-Observation condition (paired t-test: p = 0.01) and to the Self-Observation condition (paired t-test: p < 10^−5^). In the Full-Observation condition we also found higher A_ROC_ compared to the Partial-Observation condition (paired t-test: p = 0.03) and to the Self-Observation condition (paired t-test: p = 0.003). However, we found no difference between the Partial-Observation condition compared to the Self-Observation condition (paired t-test: p > 0.4, BF = 0.30) and no difference between the Baseline condition and the Full-Observation condition (paired t-test: p > 0.4, BF = 0.32).

### Questionnaire

We found no significant correlations between the emotional expressivity of participants as assessed by the Berkeley expressivity questionnaire and the relationship between the confidence of the agent and the confidence of the observer.

## DISCUSSION

In the current study we investigated the extent to which one can evaluate the confidence of others in absence of verbal communication. We also asked whether one has a privileged access in assessing confidence when performing the task directly compared to observing someone else or when observing oneself compared to observing someone else.

First, in line with the study from Patel, Fleming & Kilner (2012) (see introduction), mixed regression analyses showed that in the Full-Observation condition participants (observers) were able to judge the confidence level of agents with a good level of accuracy, indicating that verbal communication is not necessary to share confidence between members of a group. This finding is coherent with type II signal detection theory analysis, which showed that participants were not simply able to track the confidence of someone else, they could do so to the extent of being able to differentiate their correct from their incorrect responses. Indeed, the metacognitive sensitivity (A_ROC_) computed from the confidence of the observer and the performance of the agent was significantly above chance in the Full-Observation and Partial-Observation conditions.

Second, although with less accuracy, participants (observers) were as well fairly good at tracking the confidence level of agents in the Partial-Observation condition in which stimuli were not accessible. The latter finding suggest two, not necessarily exclusive, mind confidence reading mechanisms: 1. In the Full-Observation condition one could argue that participants are not simply performing the task mentally and inferring the confidence of others based on their own implicit judgements but also base their inference on the observation of the agent behaviour; 2. One could also suggest that when stimuli are not available (Partial-Observation condition) to participants, they switch to other cues. Mediation analyses revealed that response times had a stronger mediating role in the Partial-Observation than in the Full-Observation condition. One thus can conjecture that there is a shift in strategy from the Full-Observation condition to the Partial-Observation condition. Note, however, that in the Full-Observation condition, response times are still mediating part of the variance between the actual confidence of the agent and the inferred confidence by the observer. Therefore, even in the Full-Observation condition, participants do not base their inference of others confidence level entirely on their own implicit judgments. Our results rather suggest that there is a mix between task mental simulation and the observation of the agent behaviour giving an advantage in this condition in comparison to the other conditions in the assessment of others confidence.

The third important result is that there is no difference in accuracy in assessing confidence level between the Baseline condition, in which participants were performing the task, and the Full-Observation condition in which they were only observing the agent performing the task *plus* having an access to stimuli, as shown by similar metacognitive sensitivity (A_ROC_). Therefore, it seems that, at least in the current experimental design, performing the task oneself does not entail a privileged access in assessing confidence in comparison to observing someone else performing the task. In other words, a first-person perspective does not benefit participants. This could even suggest that one evaluates one’s own confidence like an external observer, that is as when one observes someone else. This finding is in line with current theoretical and modelling work (Carruthers, 2009; Fleming & Daw, 2017).

However, it might be that the potential first-person perspective advantage is obscured by the fact that task difficulty constitutes such a strong cue in assessing confidence, that is, the access participants (observers) have to stimuli in the Full-Observation condition equalizes confidence accuracy between the latter condition and the Baseline condition. In order to disentangle this point, we compared the Partial-Observation condition to the Self-Observation condition. If there is any advantage at assessing oneself *versus* someone else we should find that participants are better in the latter than in the former condition. The fourth main significant result indeed shows that the relationship between the actual confidence of the agent and the inferred confidence by the observer was stronger in the Self-Observation condition than in the Partial-Observation condition, as shown through mixed modelling analysis. It is unlikely that this results stems from a memory effect, as the Self-Observation condition was systematically performed on another day than the Baseline condition in which the video was recorded and, even if they did remember their overall level of confidence from the Baseline condition, it is inconceivable that one could remember the trial-by-trial association. Moreover, participants were not aware that they would be asked to judge their own confidence based on the video recording at the time of the Baseline condition, thus ensuring that they were not incentivized to exaggerate their behaviour so as to give their future self cues. Note however that Type 2 signal detection theory analysis showed no difference between the metacognitive sensitivity of participants in the Self-Observation condition compared to the Partial-observation condition. Further work is thus needed to explore more precisely our ability to judge ourselves from an external point of view. In addition, using mediation analyses, we found that the mediation effect was the smallest in the Self-Observation condition (in comparison to the Full- and Partial-Observation conditions), with only 26.9% of the relationship between confidences of the agent and the observer mediated by response times. This indicates that when judging themselves, participants used other cues than response times and task difficulty to judge their past confidence. Taken together, these results highlight the fact that metacognitive monitoring (of oneself or someone else) is a flexible process integrating multiple cues and that is responsive to situational demands (Reyes & Sackur, 2014; Desender, Van Opstal & Van den Bussche, 2017).

All in all, the present study demonstrates that we can successfully read the confidence of others in the absence of verbal communication or having access to the stimuli and that we can adapt the cues we rely on depending on the situation we are in. From an evolutionary perspective, this may be a crucial ability, allowing us to evaluate the confidence of our peers in various situations (Shea et al., 2014). A deeper understanding of this phenomenon may also help to shed light on several psychiatric disorders involving difficulties to read others, such as autism (Baron-Cohen, 2000; Boria et al., 2009), schizophrenia (Brüne, 2005; Walter et al., 2009) or depression (Inoue, Yamada & Kanba, 2006; Wang et al., 2008).

## METHOD

### Participants

Fifty participants were recruited to participate in the study (Mean Age = 21.3, SD = 1.8). As the experimental task involved pairs of participants, we recruited only female participants in order to avoid gender effects. All participants reported no history of hearing disorder and no psychiatric or neurological history, and participated in exchange for a monetary compensation (10€ per hour). They were naive to the purpose of the study and gave informed consent, in accordance with institutional guidelines and the Declaration of Helsinki. The study was approved by the local ethical committee of the ULB.

### Apparatus and stimuli

All experimental sounds were sinusoidal pure tones, with 5 ms rise/fall time and 44100 Hz sampling rate, generated using MATLAB (MathWorks, Natick, MA) with the Psychophysics toolbox (Brainard, 1997; Pelli, 1997; Kleiner, Brainard, Pelli 2007). Auditory stimuli used for the pitch discrimination task were chosen through pilot testing and consisted in a standard pitch sound of 500 Hz which was to be compared to 504, 508, 512, 515 or 518 Hz pitch test sounds, played for 250 ms via headphones. The reference sounds were randomly presented to the left ear or the right ear first and the test sound to the opposite ear. A fixation cross appeared prior to the sound to signal the beginning of the trial.

### Procedure

Participants were paired two by two and did not know each other. Upon arrival one participant was randomly assigned to first take the role of the agent and the other the role of the observer. They were instructed not to talk to each other.

The experiment was divided in two sessions of approximately 2 hours each. The second session took place between 24h and 48h after the first session. One session corresponded to two experimental conditions of 250 trials each (with 75 trials for 504 and 508, 50 trials for 512 and 25 trials for 515 and 518 Hz test sounds). In each condition the agent had to perform the pitch discrimination task and press the left or right arrow on the keyboard with their right hand to indicate which sound had the highest pitch. Then participants had to give their confidence in their response by pressing a key with their left hand on a separate keypad on a scale from 1 (guess) to 4 (sure). Participants then switched roles so that the agent became the observer and the observer became the agent. At each new condition they returned to their original role assignment.

In the Baseline condition (Figure 1A), participants were seated at a different desk and performed the task on their own without seeing the other participant. During this condition, they were both filmed so that their facial expression, body and hands were recorded. In the Full-Observation condition (Figure 1B) participants joined at one desk. The observer seated so that she had the same point of view as the camera in the baseline condition. The keypad on which the agent gave her confidence ratings was hidden from the observer thanks to a cardboard. Both the agent and the observer wore headphones and heard the auditory stimuli. Once the agent gave her confidence in her response the observer had to judge what she thought was the confidence of the agent by answering the same confidence scale on her own keypad. The observer could use any strategy that she wanted. Once the observer gave her response a new trial started. In the Partial-Observation condition (Figure 1C) the disposition was the same as in the Full-Observation condition except that the observer did not hear the auditory stimuli anymore and wore a sound-proof headset. However, they still had access to the response times of the agent, as a fixation cross appeared on the screen to signal the beginning of each trial. In the Self-Observation condition (Figure 1D) participants returned to their own desk as in the Baseline condition. They both took the role of the observer to judge the confidence of themselves performing the pitch discrimination task in the baseline condition by watching the recorded video without sound. One experimenter was present next to each participant to interrupt the video at each trial (a red cue was present on the screen once the participant gave her response in the Baseline condition) and waited for their confidence rating to restart the video.

The Baseline condition always took place first and the Self-Observation condition always took place last so as to avoid any memory effect in the Self-Observation condition and to ensure that all participants knew the task before judging the confidence of the agent in the following conditions. The order of the Full-Observation condition and the Partial-Observation condition was randomised over pairs of participants. At the end of the third session participants completed the Berkeley expressivity questionnaire (Gross & John, 1997) in order to assess their emotional expressivity and a debriefing.

### Data preprocessing

Data preprocessing and analyses were performed with R (2016), using the afex (Singmann et al., 2015), lme4 (Bates et al., 2014), lmerTest (Kuznetsova, Brockhoff & Christensen, 2015), BayesFactor (Morey & Rouder, 2015), ggplot2 (Wikham, 2009) and effects (Fox, 2003) packages. We discarded data from pairs of participants for which one or both participants had a mean accuracy below 55% or above 95% in at least one condition in order to be sure that both participants in a pair could perform the task. It was indeed not possible to determine individually the level of difficulty as it was critical that participants heard the same stimuli to make their confidence judgements. Twenty-three participants met this exclusion criterion and in total 15 pairs were excluded from analysis. This allowed to keep only pairs of participants who could perform the auditory task and who were not at a ceiling performance so as to be able to compute metacognitive sensitivity. Another pair of participants was discarded due to issues in the data recording during the experiment. The following analyses were thus made on 18 participants^1^Note that with all participants included in the analyses the pattern of results is virtually identical (see Supplementary Material for more details).

### Statistical analysis

In order to compare the ability of the observer to assess the confidence of the agent as well as her own confidence in the different conditions, we performed mixed models and mediation analysis. In addition, metacognitive ability was estimated through the type-II area under the Receiver Operating Curve (A_ROC_), which plots the correct response rate against the incorrect response rate at each confidence level (Kornbrot, 2006). However here, except in the Baseline condition, we use this measure in a nonconventional way, as we use the confidence of the observer and the accuracy of the agent to compute these A_ROC_. This is what we further refer to as the metacognitive ability of the observer regarding the agent.

In addition, we used within-subject repeated measures analysis of variance to test for differences in first- and second-order performances followed by paired and one sample *t*-test to determine the direction of differences. In all ANOVAs, degrees of freedom were corrected using the Greenhouse-Geisser method.

## Supporting information

Supplementary analyses

## ACKNOWLEDGEMENTS

This work was supported by an European Research Council Advanced Grant RADICAL to A.C.. The authors thank Y-Nni Tran Ngoc and Sylwia Gutowska for help with data collection.

