## Supplementary analyses for "Comparing self- and hetero-metacognition in the absence of verbal communication"

**SUPPLEMENTARY MATERIAL:**

We conducted the same analyses without removing any participants (48 subjects).

Agent performance at the first- and second-order level

First, regarding the first-order task (i.e., pitch discrimination task), a repeated-measure ANOVA showed no effect of condition on type 1 sensitivity (*d'*) ($F(1.13, 53) = 0.61$, $p > 0.4$, $\eta_p^2 = 0.01$) or criterion ($F(1.64, 77.18) = 2.45$, $p = 0.10$, $\eta_p^2 = 0.05$). However, a repeated-measures ANOVA showed differences in mean response times across the different conditions ($F(1.32, 61.98) = 28.22$, $p < 10^{-3}$, $\eta_p^2 = 0.38$). Specifically, paired *t*-tests indicated that response times were shorter in the Full-Observation (Mean = 1.41 s, SD = 0.33) and Partial-Observation (Mean = 1.37 s, SD = 0.33) conditions than in Baseline condition (Mean = 1.93 s, SD = 0.08) (Full-Observation vs. Baseline: $p < 10^{-5}$, Partial-Observation vs. Baseline: $p < 10^{-6}$), but there was no difference between the Partial- and Full-Observation condition ($p = 0.31$, BF = 0.26).

Second, with regards to the second-order task, we found no effect of condition on confidence ratings ($F(1.28, 60.39) = 0.87$, $p > 0.3$, $\eta_p^2 = 0.02$) or confidence ratings variability using standard deviation as a measure of variance ($F(1.37, 64.55) = 0.44$, $p > 0.4$, $\eta_p^2 = 0.03$).

Third, we compared the metacognitive sensitivity of the agent across the different conditions using $A_{ROC}$ as a measure of this type 2 sensitivity. A repeated-measure ANOVA showed no difference between conditions ($F(1.29, 60.41) = 1.24$, $p = 0.28$, $\eta_p^2 = 0.03$).

Observer mean confidence level across conditions

The mean confidence level of the observer did not differ between conditions ($F(1.67, 78.27) = 1.09$, $p > 0.3$, $\eta_p^2 = 0.02$) nor did the confidence variability ($F(1.60, 75.04) = 1.13$, $p > 0.3$, $\eta_p^2 = 0.02$).

Observer ability to read agent confidence

Before comparing the relation between the observer and the agent confidence between the different conditions, we checked whether this relation was significant within each condition. For each condition, we fitted a linear mixed-effects model of the confidence of the observer, with confidence of the agent as fixed effect and intercept for participants as random effect. Results confirm that the observer was indeed capable of judging the confidence of the agent in each condition (Full-Observation: estimate = 0.20, t = 23.89, $p < 10^{-3}$; Partial-Observation: estimate = 0.17, t = 20.68, $p < 10^{-3}$; self-observation: estimate = 0.15, t = 18.44, $p < 10^{-3}$).

Then, we compared the relations between the agent's and observer's confidences between these different conditions. We fitted a linear mixed-effects model of the confidence of the observer, with confidence of the agent and condition (Full-Observation, Partial-Observation and Self-Observation) as fixed effects and intercept for participants as random effect.

| | Estimate | SE | T | *p* |
|---|---|---|---|---|
| Intercept | 2.26 | 0.06 | 40.43 | <0.001 |
| Confidence of the doer | 0.19 | 0.01 | 23.90 | <0.001 |
| Partial-Observation condition | 0.11 | 0.03 | 3.37 | <0.001 |
| Self-Observation condition | 0.05 | 0.03 | 1.63 | 0.1 |
| Confidence of the doer: Partial-Observation condition | -0.04 | 0.01 | -3.56 | <0.001 |
| Confidence of the doer: Self-Observation condition | -0.04 | 0.01 | -3.55 | <0.001 |

**Number of participants: 48**
**Number of observations: 36 000**

Table S1. Regression coefficients for the linear mixed-effects model of the confidence of the observer in the three conditions.

The first row in **Table S1** (intercept) estimates the average confidence of the observer in the Full-Observation condition for the lowest scale rating of the confidence of the agent. The observer had a significantly higher confidence than the agent when the latter reported guessing (estimate = 2.26, t = 40.43, $p < 10^{-3}$).

The second row estimates the regression slope between the confidence of the observer and the confidence of the agent in the Full-Observation condition, and shows that this relation is statistically significant (estimate = 0.19, t = 23.90, $p < 10^{-3}$), indicating that the observer can track the confidence of the agent.

The third and fourth row of the model show that the confidence of the observer for the lowest scale rating of the agent was not significantly different between the Self-Observation condition and Full-Observation condition (estimate = 0.05, t = 1.63, p = 0.1), but was significantly higher in the Partial-Observation compared to the Full-Observation condition (estimate = 0.11, t = 3.37, $p < 10^{-3}$).

The fifth and the sixth row indicate that the relation between the confidence of the observer and the confidence of the agent was smaller in the Partial-Observation compared to the Full-Observation condition (estimate = - 0.04, t = - 3.56, $p < 10^{-3}$) and in the Self-Observation compared to the Full-Observation condition (estimate = - 0.04, t = - 3.55, $p < 10^{-3}$).

Another linear mixed-effects model comparing only the Self-Observation condition (in which participants were judging their own performances in the baseline condition by means of video recording) to the Partial-Observation condition revealed no difference in intercepts or regression slopes between the confidence of the observer and the confidence of the agent in the Self-Observation condition compared to the Partial-Observation condition.

### Do observers read agents' confidence from their response times

We used causal mediation analyses to test whether the effect of the confidence of the agent on the confidence of the observer was mediated by the response times of the agent. In each condition, a mediator mixed model was first fitted to predict the response times of the agent by the agent's confidence. Then, an outcome mixed model was fitted to predict the confidence of the observer by the response times and the confidence of the agent. The mediation analysis was performed with these two models (using the mediation package; Tingley, Yamamoto, Hirose, Keele, & Imai, 2014) in order to test whether the influence of the confidence of the agent on the confidence of the observer was mediated by the response times of the agent.

In the Full-Observation condition the mediation analysis showed that from the total effect of the confidence of the agent on the confidence of the observer ($\beta = 0.204$, 95% CI = [0.186, 0.222], $p < .001$), there was 41.6% that was mediated by the response times of the agent ($\beta = 0.085$ 95% CI = [0.078, 0.093], $p < .001$). In the Partial-Observation condition, from the total effect of the confidence of the agent on the confidence of the observer ($\beta = 0.172$, 95% CI = [0.155, 0.187], $p < .001$), there was 49.0% that was mediated by the response times of the agent ($\beta = 0.084$, 95% CI = [0.077, 0.092], $p < .001$). In the Self-Observation condition, from the total effect of the confidence of the agent on the confidence of the observer ($\beta = 0.151$, 95% CI = [0.135, 0.168], $p < .001$), there was 28.4% that was mediated by the response times of the agent ($\beta = 0.043$, 95% CI = [0.036, 0.049], $p < .001$).

### Type-II signal detection theory

A one-way one sample t-test showed that the $A_{ROC}$ of participants judging themselves in the Baseline condition were significantly higher than 0.5 ($p < 10^{-9}$). In the Full-, Partial- and Self-Observation conditions the $A_{ROC}$ were also significantly higher than 0.5 (Full-Observation condition: $p < 10^{-8}$; Partial-Observation condition: $p < 10^{-3}$, Self-Observation condition: $p < 0.01$), suggesting that the metacognitive ability of the observer regarding the agent (or herself through video recording in the Self-Observation condition) was also higher than chance.

Analysis of variance revealed a significant difference between conditions ($F(2.49, 116.90) = 15.25$, $p < 10^{-4}$, $\eta_p^2 = 0.24$), with higher $A_{ROC}$ in the Baseline condition compared to the Partial-Observation condition (paired t-test: $p < 10^{-3}$) and to the Self-Observation condition (paired t-test: $p < 10^{-7}$). In the Full-Observation condition we also found higher $A_{ROC}$ compared to the Partial-Observation condition (paired t-test: $p < 10^{-4}$) and to the Self-Observation condition (paired t-test: $p < 10^{-4}$). However, we found no difference between the Partial-Observation condition compared to the Self-Observation condition (paired t-test: $p > 0.4$, BF = 0.18) and no difference between the Baseline condition and the Full-Observation condition (paired t-test: $p > 0.4$, BF = 0.17).

<u>Questionnaire</u>

We found no significant correlations between the emotional expressivity of participants as assessed by the Berkeley expressivity questionnaire and the relationship between the confidence of the agent and the confidence of the observer.
